# Role of Nonneutralizing Antibodies and Fc Effector Functions in Inhibiting SARS-CoV-2 Infection

**DOI:** 10.64898/2026.04.08.717316

**Authors:** Haiyan Sun, Adrian Esqueda, Herta Steinkelner, Qiang Chen

**Affiliations:** The Biodesign Institute and School of Life Sciences, Arizona State University, Tempe, Arizona, United States of America; Department of Applied Genetics and Cell Biology, University of Natural Resources and Life Sciences, 1180 Vienna, Austria

**Keywords:** Fc effector function, Monoclonal antibody (mAb), SARS-CoV-2, Antibody-dependent cellular cytotoxicity (ADCC), Glycosylation, Nonneutralizing antibody, antibody-dependent cell-mediated virus inhibition (ADCVI), Plant-made antibody, Plant-made pharmaceutical

## Abstract

Neutralizing monoclonal antibodies (mAbs) are a key component of antiviral therapeutics against SARS-CoV-2; however, the contribution of Fc-mediated effector functions remains underexplored. Here, we compare the antiviral activities of the neutralizing and non-neutralizing mAbs CB6 and CR3022, respectively. The Fc regions of both plant-produced mAbs carried nonfucosylated, non-galactosylated complex glycans (pCB6 and pCR3022), and CR3022 was also produced with mammalian-typical galactosylated, fucosylated glycans (mCR3022). pCR3022 exhibited markedly enhanced antibody-dependent cellular cytotoxicity (ADCC) and antibody-dependent cell-mediated virus inhibition (ADCVI) compared to mCR3022, indicating a significant impact of Fc glycosylation on antiviral activity despite the lack of neutralization. pCB6 exhibited potent neutralization while further enhancing virus clearance through synergistic Fc effector activity. Our findings suggest that Fc-mediated mechanisms, especially ADCC and ADCVI, can contribute substantially to viral control and may be particularly valuable against immune-evasive variants. These results advance our understanding of the functional roles that non-neutralizing antibodies can play in SARS-CoV-2 infection and highlight the potential of Fc glycoengineering to modulate the antiviral efficacy of both neutralizing and non-neutralizing mAbs.

## Introduction

The COVID-19 pandemic, caused by SARS-CoV-2, has led to unprecedented global health and socioeconomic disruption since its emergence in late 2019. With over 778 million confirmed cases and over 7 million deaths reported worldwide as of 2025 [1], the pandemic has placed immense pressure on public health systems and driven an urgent need for effective therapeutics.

Among the most impactful biomedical interventions developed during the pandemic are monoclonal antibodies (mAbs). Under the U.S. Emergency Use Authorization (EUA) program, multiple SARS-CoV-2-targeting mAbs — including bamlanivimab, casirivimab/imdevimab, tixagevimab/cilgavimab, and sotrovimab — were rapidly deployed [2]. These mAbs offer high specificity and potent neutralization capacity. However, several drawbacks limit their widespread and sustained use: they are expensive to manufacture, especially in mammalian cell culture-based systems; they are vulnerable to loss of efficacy due to viral escape mutations; and their neutralization breadth is often narrow, requiring constant updates to keep pace with emerging variants [2,3].

In addition to neutralizing antibodies, infection and vaccination also elicit non-neutralizing antibodies. While the protective roles of neutralizing antibodies are well-established, the contributions of non-neutralizing antibodies remain less clearly defined [4]. Some studies suggest that they may provide protection through mechanisms independent of neutralization [5,6], while others raise concerns about their potential to exacerbate disease through antibody-dependent enhancement (ADE) [7,8]. Understanding the role of non-neutralizing antibodies in viral immunity is essential for the rational design of vaccines and antibody-based therapeutics. In addition, the contribution of Fc-mediated effector functions to either antiviral activity or disease enhancement by non-neutralizing antibodies, as well as their ability to affect the effectiveness of neutralizing antibodies, remains incompletely understood.

Plant-based expression platforms have emerged as powerful tools for recombinant protein production [9] and have played a meaningful role in advancing biomedical countermeasures during the COVID-19 pandemic [10,11]. One notable success is the development of a virus-like particle (VLP) vaccine displaying a modified SARS-CoV-2 spike protein, produced in plants [12]. This plant-derived vaccine demonstrated favorable safety and efficacy profiles in clinical trials and has received regulatory approval for human use in Canada, marking a major milestone in plant-based biopharmaceuticals [12]. Plants also offer a promising and increasingly recognized alternative to conventional mammalian cell culture systems for the development and production of mAbs [13,14]. Mammalian platforms, while effective, are often constrained by high production costs and the risk of contamination with adventitious human pathogens. In contrast, plant systems provide a scalable, cost-effective, and pathogen-free alternative that is particularly well suited for rapid response and flexible engineering [15–17]. One key advantage of plant expression systems is their amenability to N-glycan modification, which enables the generation of mAbs with tailored glycosylation profiles [18,19]. This capacity for glycoengineering will reduce glycan heterogeneity and may allow the development of antibodies with enhanced Fc-mediated functions. Furthermore, plant-based transient expression via agroinfiltration enables rapid and high-yield production of mAbs in a matter of days, accelerating the development timeline compared to traditional platforms [20–23].

A growing body of evidence supports the viability of plant-produced mAbs as functionally superior alternatives to their mammalian-derived counterparts. Multiple plant-derived antibodies have demonstrated strong performance in preclinical models, including high antigen specificity, functional activity, and consistent glycosylation [17,24–28]. Collectively, these attributes underscore the potential of plant systems not only for scalable antibody manufacturing but also for developing next-generation antibody variants with optimized therapeutic properties.

Here, we used CR3022, a non-neutralizing antibody targeting the receptor-binding domain (RBD) of SARS-CoV-2, and CB6, a neutralizing mAb, as models to investigate the impact of Fc glycan engineering in antiviral activities. Glycoengineered *Nicotiana benthamiana* plants were used to produce CR3022 and CB6 with homogeneous, fucose free glycans (GnGn) [29]. Both CR3022 and CB6 produced in glycoengineered plants exhibited the expected homogeneous GnGn glycosylation profile and retained antigen-binding specificity comparable to their mammalian cell-produced counterparts reported in previous studies. The neutralization potency of plant-produced CB6 was also consistent with previously reported results. Plant-derived CR3022 exhibited enhanced Fc effector activity, including antibody-dependent cellular cytotoxicity (ADCC) and antibody-dependent cell-mediated virus inhibition (ADCVI). Notably, such Fc-dependent mechanisms enabled plant-derived CR3022 to mediate significant viral clearance despite lacking neutralizing activity. In parallel, plant-produced CB6 demonstrated a synergistic enhancement of viral elimination when both neutralization and Fc effector functions were engaged.

## Material and methods

### Production and purification of mAbs from *N. benthamiana* plants

The genes encoding the variable regions of the heavy chain (HC) and light chain (LC) of CR3022 [30] were synthesized and cloned into vectors containing the coding sequences for the human IgG1 HC constant region and kappa LC constant region, respectively [11]. These HC and LC genes were subsequently cloned into a geminiviral-based plant expression vector and introduced into *Agrobacterium tumefaciens* as previously described [25]. The *A. tumefaciens* strain carrying the CR3022 HC and LC construct was then used to infiltrate the leaves of 6-week-old ΔXF *N. benthamiana* plants, following established procedures [22,23,31]. CR3022 mAb was extracted and purified from plants following protocols previously described by our group [11,26], In brief, leaves were collected at 7 days post infiltration (DPI), homogenized in extraction buffer (50 mM Tris-HCl pH7.5, 150 mM NaCl), and the total leaf soluble protein extracts were clarified by centrifugation at 15,000 x *g* for 30 min at 4°C. The resulting supernatant was filtered through a 0.2-micron filter and the mAb was purified via a Protein A-based protocol previously described [32]. Purified mAbs were analyzed by SDS-PAGE under reducing conditions using 10% gradient acrylamide gels (Bio-Rad). Gels were stained with Coomassie Brilliant Blue R-250 to visualize total protein content, as previously described [25]. The expression, purification and characterization of the CB6 mAb, a neutralizing antibody produced in plants have been reported in prior work [33]. Mammalian cell-produced CR3022 was purchased commercially (Abcam).

### Antigen-Specific Binding ELISA

The binding specificity of mAbs to the SARS-CoV-2 RBD was assessed by ELISA, following a previously described protocol [33]. Briefly, 96-well ELISA plates were coated overnight at 4□°C with 2□µg/mL of recombinant RBD protein (WA1/2020 strain) in phosphate-buffered saline (PBS). Plates were then blocked with 5% nonfat milk in 1× PBS containing 0.05% Tween-20 (PBST) for 1□h at room temperature. After blocking, serial dilutions of mAbs were prepared in 5% milk in PBST and incubated on the plates for 1□h at 37□°C. Following three washes with PBST, bound antibodies were detected using horseradish peroxidase (HRP)-conjugated goat anti-human IgG secondary antibody (Southern Biotech). TMB substrate (SeraCare Life Sciences) was added, and plates were developed for 5 min before the reaction was stopped with 1□M H_₂_SO_₄_. Absorbance was measured at 450□nm using a microplate reader. GraphPad Prism version 10.2.2 was used to generate binding curves and calculate approximate dissociation constants (KD) using a one-site specific binding model.

### Analysis of N-linked glycosylation of mAbs

The N-glycosylation profiles of the purified mAbs were assessed using mass spectrometry (MS) based on previously described methods [34,35]. Briefly, approximately 6 µg of purified antibodies were subjected to trypsin digestion and analyzed using a liquid chromatography–electrospray ionization–mass spectrometry (LC-ESI-MS) system (Thermo Orbitrap Exploris 480). Glycopeptides were identified by their characteristic peak patterns, which represented the peptide backbone with attached N-glycans differing in the number of HexNAc units, hexose, deoxyhexose, and pentose residues. Glycopeptide identification was conducted using FreeStyle 1.8 (Thermo), with deconvolution performed via the extract function. The relative molar ratios of the glycoforms were estimated based on peak heights. Glycan nomenclature adhered to the guidelines as described in [36].

### Immunofluorescence Staining

Immunofluorescence staining was performed as previously described [37]. In brief, Vero E6 cells (5 × 10□) were seeded into each well of a 12-well tissue culture plate. The following day, cells were infected with SARS-CoV-2 (USA-WA1/2020 strain) at a multiplicity of infection (MOI) of 1. The virus was diluted in serum-free DMEM and added to the monolayers for 1 h at 37□°C. After incubation, the inoculum was removed, cells were washed three times with PBS, and fresh DMEM supplemented with 2% fetal bovine serum (FBS) was added. Infected cells were then incubated at 37□°C for an additional 24 h. Cells were fixed with 4% paraformaldehyde for 15 min at room temperature and permeabilized with 0.1% Triton X-100 in PBS. For staining, cells were incubated with a rabbit anti-nucleocapsid (N) monoclonal antibody (Cell Signaling Technology; 1:2000 dilution) and plant-produced CR3022 (5□µg/mL). Secondary antibodies included Alexa Fluor 647-conjugated goat anti-rabbit IgG (H+L) (Thermo Fisher Scientific) and Alexa Fluor 488-conjugated goat anti-human kappa light chain antibody (SouthernBiotech). Finally, cells were stained with DAPI (Lifetech) for nucleus visualization. Images were captured using a fluorescence microscope.

### Neutralization against SARS-CoV-2

The neutralizing activity of CB6 against SARS-CoV-2-delta3a/7b [38] was assessed using a focus-forming assay (FFA), as previously described [11]. Briefly, 25,000 Vero-hACE2-TMPRSS2 cells were seeded in 96-well flat-bottom clear plates in 100 µL of DMEM supplemented with 10% FBS one day prior to infection. On the day of the assay, serial dilutions of CB6 were prepared, and viral stocks were diluted to 2000 plaque-forming units (PFU). The diluted CB6 was incubated with the virus for 1 h at 37 °C, then the mixture was added to the cell monolayers and incubated for an additional hour at 37 °C. All antibody and virus dilutions were prepared in DMEM containing 2% FBS. Following infection, a 100 µL overlay of MEM containing methylcellulose was applied to each well. Infected cells were incubated for 24 h at 37 °C. After incubation, the overlay was removed, and cells were fixed with 4% paraformaldehyde. Cells were washed six times with PBS containing 0.1% saponin and 0.1% bovine serum albumin (BSA), then stained with a rabbit anti-N monoclonal antibody (Cell Signaling Technology). Detection was performed using goat anti-rabbit IgG-HRP (Sigma). Foci were developed using KPL TrueBlue substrate (SeraCare Life Sciences Inc.) and quantified using an AID Spot Reader.

Percent neutralization was calculated as follows: (mean number of foci in virus-only control wells − number of foci in antibody-treated wells) / mean number of foci in virus-only control wells. Data were analyzed using GraphPad Prism 10.2.2, and EC_₅₀_, EC_₂₅_, and EC_₇₅_ values were calculated. All experiments were independently repeated at least twice with triplicate samples

### Antibody-Dependent Cellular Cytotoxicity Reporter Assay

The ADCC reporter bioassay kit (Promega) was used to evaluate antibody-mediated cellular cytotoxicity, following the manufacturer’s protocol. Target cells stably expressing the SARS-CoV-2 spike protein (InvivoGen) were seeded at 20,000 cells per well in white 96-well tissue culture plates one day prior to the assay. Serial 1:5 dilutions of mAbs were prepared in ADCC assay buffer (RPMI supplemented with 4% low IgG serum). Effector cells (Jurkat cells expressing Fc gamma receptor IIIa (FcγRIIIa)) were added at a density of 75,000 cells per well, as specified by the Promega protocol. After the mAb and effector cells were added to the target cell plates, they were incubated for 19–20 h at 37□°C to allow for effector-target interaction. After incubation, the luminescent substrate was added, and luminescence was measured using a SpectraMax M5 plate reader (Molecular Devices). Background signal from without mAb added was subtracted from the measurement of each well. The data were normalized and EC_50_ values were calculated using GraphPad Prism 10.2.2. Data represent combined results from a minimum of two independent experiments, each performed in technical triplicate.

### Antibody-Dependent Cell-Mediated Virus Inhibition Assay

Human peripheral blood mononuclear cells (PBMCs) were isolated from buffy coats obtained from BioIVT, following previously described protocols [10]. Frozen PBMCs were recovered overnight at 37□°C in RPMI medium supplemented with 10% FBS and 50 U/mL recombinant human IL-2 one day prior to the assay. Vero-hACE2-TMPRSS2 cells were seeded at a density of 1.5 × 10□ cells per well in 24-well tissue culture plates and incubated overnight. The SARS-CoV-2-delta3a/7b virus stock was diluted in RPMI containing 2% FBS to a final concentration of 500 focus-forming units (FFU) per well. On the day of infection, culture media were removed, and the diluted virus was added to the cells for 40–45 min at 37□°C. Following infection, CR3022 was diluted to 100□µg/mL, and CB6 was diluted to concentrations at 10-fold of the calculated EC_₂₅_, EC_₅₀_, or EC_₇₅_ values as determined by the FFA above, with RPMI supplemented with 2% FBS. A total of 50□µL of the diluted mAbs was added to each well. Plates were then incubated for 15–20 min at 37□°C prior to the addition of PBMCs. PBMCs were counted, pelleted by centrifugation at 400 × g for 5 min, and resuspended in RPMI + 2% FBS at a concentration of approximately 7.5 × 10□ cells/mL. A total of 200 µL of the PBMC suspension was added to each well to achieve an effector-to-target (E:T) cell ratio of 10:1. After 24 h of co-culture, supernatants were harvested, centrifuged at 400 × g for 5 min to remove cellular debris, aliquoted, and stored at −80□°C. Viral titers in the supernatants were quantified by FFA using serial dilutions as described above. Data were normalized by setting the virus titer from the “virus only” control supernatants to 100%. Each condition was tested in at least two independent experiments with technical triplicates, and normalized values were combined for statistical analysis using GraphPad Prism 10.2.2.

### Synergy Analysis

To assess potential synergy between mAb neutralization and effector cell-mediated viral inhibition, the mAb was tested at a concentration corresponding to its EC_₇₅_ value. Experiments were performed using three conditions: neutralizing mAb alone, PBMCs + non-neutralizing mAb at equivalent concentration, and neutralizing mAb in combination with PBMCs. The observed viral titer reduction was quantified using the ADCVI assay, as described above. The resulting data were imported into the SynergyFinder platform [39] and evaluated using four standard drug interaction models: Highest Single Agent (HSA) [40], Loewe additivity (LOEWE) [41], Bliss independence (BLISS) [42], and Zero Interaction Potency (ZIP) [43]. For each condition and concentration, the predicted percentage of viral inhibition was calculated and compared to the experimentally observed values. Synergy was defined as occurring when the actual viral inhibition exceeded the predicted effect across all models.

### Statistical analyses

Statistical analyses were performed using GraphPad Prism version 10.2.2. T-test was used to compare ADCC activity induced by different mAb glycovariants and compare virus titers between treatment groups involving different mAb glycovariants, with or without PBMCs, in the ADCVI assay. A p-value of < 0.05 was considered statistically significant.

## Results

### Characterization of plant-produced CR3022 mAb

CR3022 mAb was produced in six-week-old glycoengineered ΔXF *N. benthamiana* plants [29] via transient expression. SDS-PAGE analysis confirmed the presence of both HC and LC at the expected molecular weights (**Figure S1**). The antibody was purified using protein A chromatography and exhibited a degree of homogeneity comparable to that of a pharmaceutical-grade isotype IgG produced in mammalian cell culture (**Figure S1**).

The antigen-binding activity of plant-produced CR3022 (pCR3022) was evaluated by immunofluorescence microscopy and ELISA. Immunofluorescence microscopy demonstrated that pCR3022 specifically bound to SARS-CoV-2-infected Vero cells (**Figure S2, Panel A**), with no detectable staining in uninfected controls (**Figure S2, Panel D**). Co-staining with an anti-nucleocapsid monoclonal antibody (Cell Signaling Technology) confirmed that pCR3022-positive cells were indeed SARS-CoV-2-infected, as both antibodies labeled the same cell population (**Figures S2, Panel A, B, and C**). ELISA analysis further demonstrated that pCR3022 binds to the SARS-CoV-2 RBD in a specific and dose-dependent manner (**Figure S3**). The dissociation constant (KD) was calculated to be 0.31 nM, consistent with previously reported affinities for mammalian cell-produced CR3022 (mCR3022) [30], indicating high-affinity binding to the RBD. As expected, the IgG isotype control showed no detectable binding (**Figure S3**). These results confirm that pCR3022 retains proper structural integrity and antigen specificity, comparable to mCR3022.

### N-linked glycan profiles of mAbs produced in different expression systems

Given that glycan composition plays a key role in modulating IgG Fc-mediated effector functions [44], we performed N-glycan profiling of CR3022 and CB6 expressed in different host cells using LC-ESI-MS. Analysis revealed that pCR3022 and plant-produced CB6 (pCB6) expressed in ΔXF glycoengineered *N. benthamiana* displayed GlcNAc-terminated complex N-glycans - predominantly the GnGn structure – and lacked xylose and fucose residues (**Figure 1**). Both antibodies demonstrated high glycan uniformity, with a single predominant glycoform comprising 81.8% of total N-glycans for pCR3022 and 92.7% for pCB6. A small proportion of high-mannose structures was also detected. In contrast, mCR3022 produced in mammalian cells exhibited a more heterogeneous glycosylation profile of four glycoforms. Among these, two dominant species corresponded to core α1,6-fucosylated structures, either with or without terminal β1,4-galactose residues (GnGnF_6_ and AAF_6_) (**Figure 1**).

**Figure 1.**
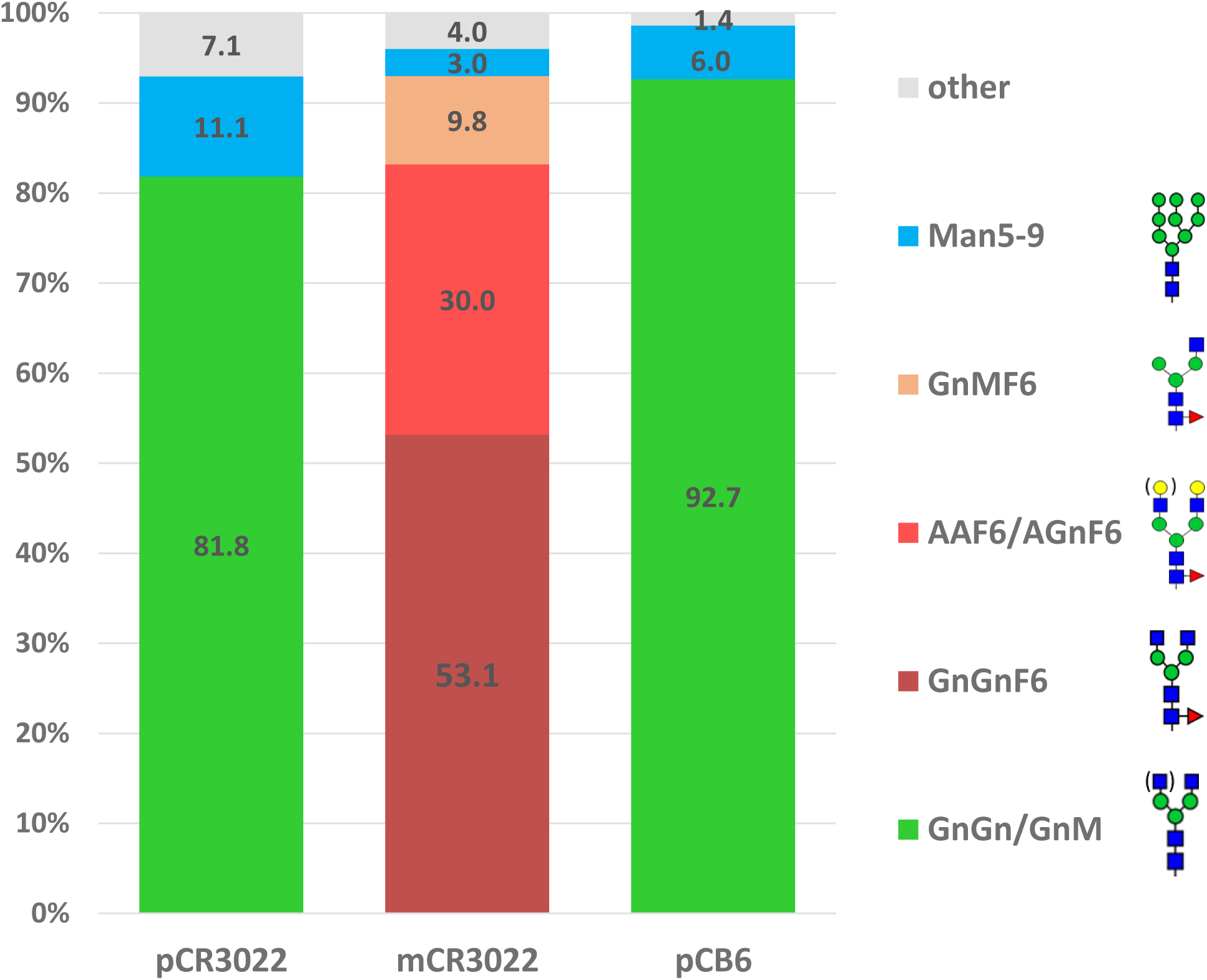
N-glycan profiles of mAbs produced in glycoengineered plants and mammalian cells. Bar graphs show the relative abundance of N-linked glycoforms determined by LC-ESI-MS analysis of CR3022 and CB6 mAbs produced in ΔXF glycoengineered *N. benthamiana* plants (pCR3022 and pCB6) and in CHO cells (mCR3022). Green bars: non-fucosylated complex GlcNAc-terminating N-glycans (predominantly GnGn structures); blue bars: mannosidic glycans (Man5–Man9); the red bar: fucosylated complex glycans with terminal galactose; the orange bar: fucosylated complex GlcNAc-terminating glycans; and gray bars: all glycoforms detected at less than 3% relative abundance. Glycan structures are labeled according to [36]

### ADCC Activity of CR3022 Glycovariants

The ADCC activity of p- and mCR3022 was evaluated using a luciferase-based reporter assay. Both mAb variants induced luciferase activity in effector cells in the presence of target cells, indicating ADCC activation (**Figure 2**). In contrast, the isotype control IgG did not elicit any luciferase signal (**Figure 2**). Notably, pCR3022 demonstrated significantly greater ADCC activity compared to mCR3022, with EC_₅₀_ values of 0.32 µg/mL and 27.93 µg/mL, respectively (*p* = 0.027) (**Figure 2**).

**Figure 2.**
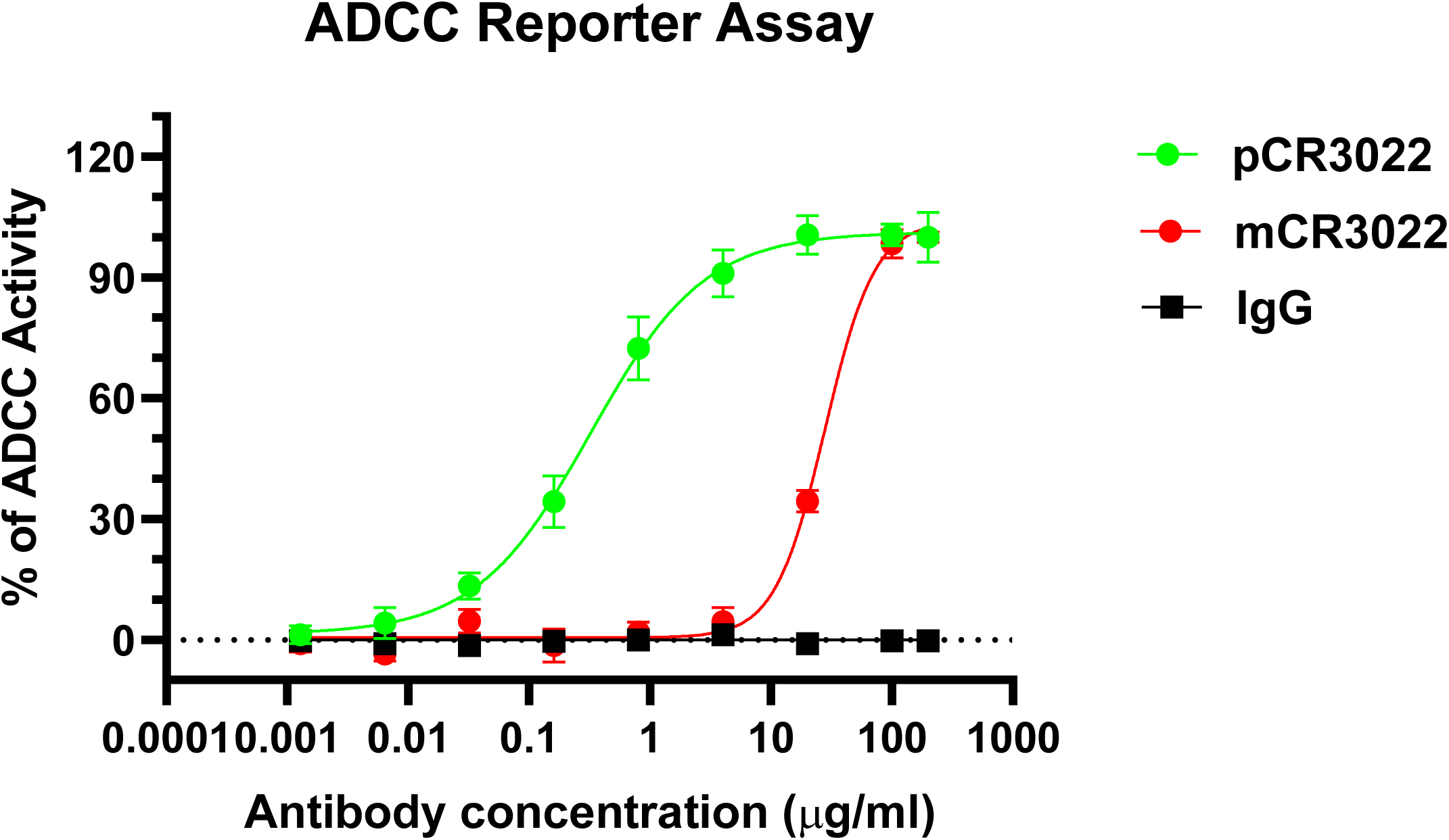
ADCC activity of CR3022 glycovariants against SARS-CoV-2. Antibody-dependent cellular cytotoxicity was assessed using the Promega ADCC Reporter Bioassay. Serial dilutions of CR3022 glycovariants or an IgG isotype control were incubated with engineered Jurkat effector cells expressing FcγRIIIa and target cells stably expressing the SARS-CoV-2 spike protein. Antibody-mediated effector-target engagement was quantified by measuring luciferase activity. Data were analyzed and EC_₅₀_ values calculated using GraphPad Prism (version 10.2.2). Results shown represent data (Mean ± SD) from at least three independent experiments, each performed in technical triplicates.

### ADCC activity enables elimination of infectious SARS-CoV-2 by a non-neutralizing antibody

We further assessed whether the ADCC activity of CR3022 glycovariants, a non-neutralizing antibody, could mediate the elimination of infectious SARS-CoV-2 using an ADCVI assay. As anticipated, treatment with either pCR3022 or mCR3022 alone did not significantly reduce viral titers compared to the virus-only control (**Figure 3**; Virus + pCR3022 vs. Virus only: *p* = 0.082; Virus + mCR3022 vs. Virus only: *p* = 0.069; Virus + pCR3022 vs. Virus + mCR3022: *p* = 0.922). Treatment with effector cells (PBMCs) alone had a minor effect on viral titers (Virus + PBMC vs. Virus only: *p* = 0.017).

**Figure 3.**
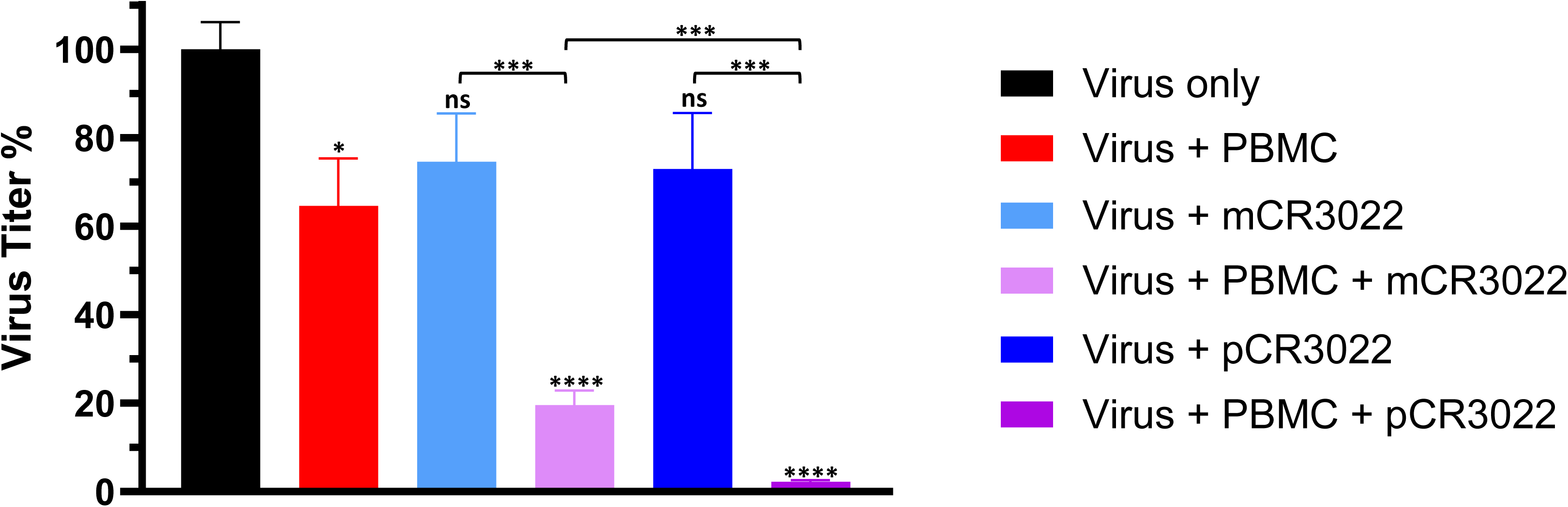
ADCVI activity of CR3022 glycovariants. Equal amount of CR3022 glycovariants were added to Vero-hACE2-TMPRSS2 cells infected with SARS-CoV-2-delta3a/7b. Buffer alone served as the “Virus only” negative control. Following antibody incubation, PBMCs were added at an E:T ratio of 10:1. In parallel, PBMC dilution buffer was used as a control for this step. After co-culture, viral titers in supernatants were quantified using an FFA. Data were normalized to the “Virus only” control, which was set to 100%. Results are presented as mean ± SD from at least two independent experiments, each performed in technical triplicates. Data analysis was conducted using GraphPad Prism 10.2.2. Asterisks and "ns" directly above each bar indicate the statistical difference between the indicated treatments (CR3022 variants, PBMCs, or both) and the “Virus only” control. ****, ***, *, and ns denote p values < 0.0001, < 0.00071, = 0.017, and > 0.05, respectively.

In contrast, co-incubation of infected target cells with both antibody and PBMCs resulted in a significant reduction in viral titers compared to all control conditions (**Figure 3**). Specifically, viral titers were significantly decreased when pCR3022 or mCR3022 was combined with PBMCs compared to Virus only (*p* < 0.0001 for both), pCR3022 or mCR3022 alone (pCR3022: *p* = 0.0002; mCR3022: *p* = 0.0007), or PBMCs alone (pCR3022: *p* = 0.0002; mCR3022: *p* = 0.0025). Notably, pCR3022 demonstrated significantly greater virus-eliminating potency than mCR3022 in the presence of PBMCs (Virus + PBMC + pCR3022 vs. Virus + PBMC + mCR3022: *p* = 0.0004).

These findings indicate that Fc-mediated effector functions can enable even non-neutralizing antibodies to eliminate infectious SARS-CoV-2. Furthermore, Fc glycoengineering, in this case the GnGn glycoform, enhances ADCC activity and improves antiviral efficacy in eliminating live viruses.

### Fc-Mediated Effector Functions Synergize with Neutralization to Eliminate SARS-CoV-2

Building on these findings, we next investigated whether Fc-mediated effector functions could similarly enhance the antiviral activity of neutralizing antibodies. We selected pCB6, a glycoengineered version of the neutralizing antibody CB6, for evaluation in both neutralization and ADCVI assays.

Neutralization assays confirmed that glycoengineering did not impair the neutralizing function of CB6, as pCB6 exhibited an EC_₅₀_ of 0.97 nM — closely matching previously reported EC_₅₀_ value of 0.93 nM for CB6 [33] (**Figure S4**). To assess Fc effector function-driven antiviral effects, we evaluated pCB6 in the ADCVI assay at concentrations corresponding to its EC_₂₅_, EC_₅₀_, and EC_₇₅_. In the absence of PBMCs, pCB6 reduced viral titers to 72.61%, 55.11%, and 27.37% of the control (defined as 100%) at EC_₂₅_, EC_₅₀_, and EC_₇₅_, respectively (**Figure 4**). These reduction percentages correspond well to the antibody concentrations used, consistent with neutralization being the predominant mechanism of viral elimination in the absence of Fc effector cells.

**Figure 4.**
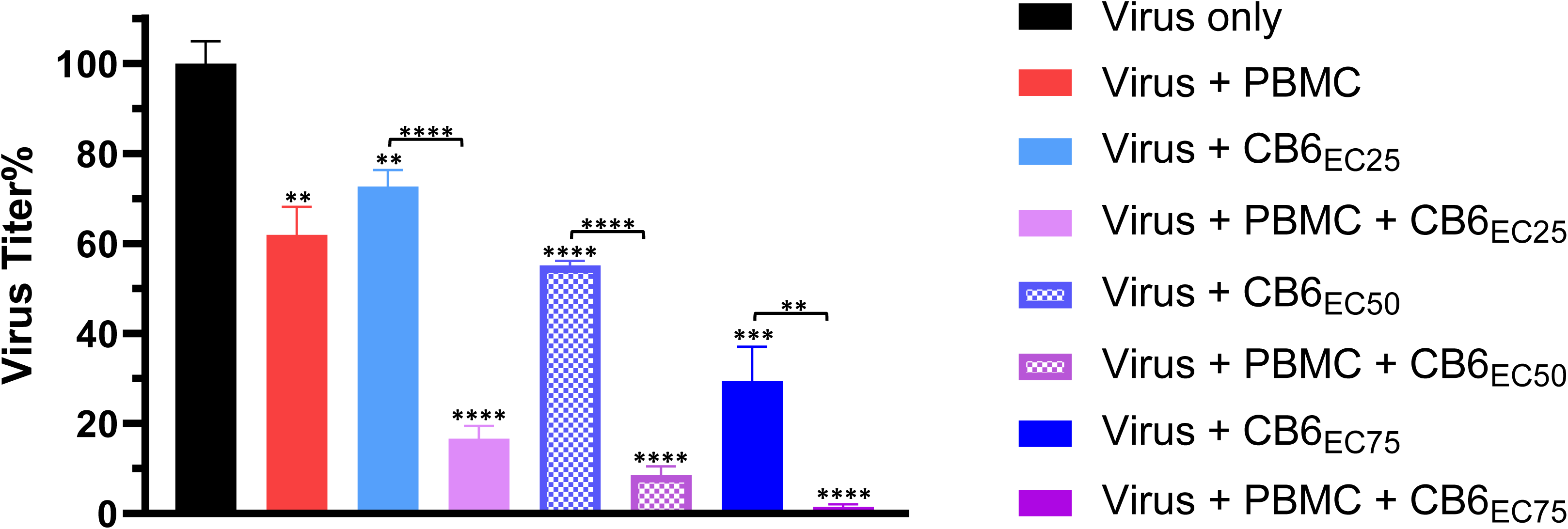
ADCVI activity of pCB6 at different concentrations. pCB6 was applied to SARS-CoV-2-delta3a/7b–infected Vero-hACE2-TMPRSS2 cells at concentrations corresponding to its EC_₂₅_, EC_₅₀_, and EC_₇₅_ values. A buffer-only condition served as the “Virus only” negative control. After antibody incubation, PBMCs were introduced at an E:T ratio of 10:1. PBMC dilution buffer was included as a secondary negative control. Following co-culture, viral titers in the supernatant were measured using an FFA. Data are shown as mean ± SD from a minimum of two independent experiments, each conducted in technical triplicates. Statistical analyses were performed using GraphPad Prism (version 10.2.2). Asterisks shown above each bar indicate statistical comparisons between the indicated conditions and the “virus only” control. ****, ***, and ** indicate p values < 0.0001, = 0.0002, and < 0.0035, respectively.

Importantly, co-incubation with PBMCs further and significantly enhanced viral clearance across all tested concentrations. Specifically, viral titers were significantly lower when pCB6 was combined with PBMCs compared to either antibody alone (EC_₂₅_: p < 0.0001; EC_₅₀_: p < 0.0001; EC_₇₅_: p = 0.0034) or PBMCs alone (EC_₂₅_: p = 0.0003; EC_₅₀_: p = 0.0001; EC_₇₅_: p < 0.0001) (**Figure 4**). To determine whether the enhanced viral reduction observed with the combination of pCB6 and PBMCs was more than additive, we conducted a synergy analysis using the SynergyFinder platform [39]. Viral reduction data from pCB6 alone (neutralization only control), PBMCs + pCR3022 (ADCC only control), and the combination of pCB6 + PBMCs were analyzed using four established reference models: HSA [40], LOEWE [41], BLISS [42], and ZIP[43]. These models estimate the expected antiviral effect under the assumption that there is no synergistic interaction between neutralization and Fc effector function-mediated reduction. Across all models, the observed viral reduction from the combination of pCB6 and PBMCs consistently exceeded the predicted additive effects (**Table S1**), indicating a synergistic interaction between the two mechanisms of viral elimination.

## Discussion

The devastating consequences of the COVID-19 pandemic have underscored the urgent need for effective therapeutics, particularly for individuals who are allergic to vaccines or immunocompromised patients who do not mount sufficient responses to vaccination [2]. While the roles of neutralizing antibodies in preventing and treating SARS-CoV-2 infection are well established, the functional significance of non-neutralizing antibodies remains poorly understood [8]. Moreover, the influence of Fc-mediated effector functions—whether protective or potentially pathogenic—on the activities of both neutralizing and non-neutralizing antibodies is not fully elucidated [7]. In this study, we investigated how Fc glycan composition affects the antiviral function of neutralizing and non-neutralizing mAbs. We show that plant-derived mAbs carried homogeneous mammalian-type GnGn glycoforms, in contrast to the more heterogeneous glycosylation profiles typically observed in antibodies produced in mammalian cell lines. This uniform glycosylation allowed us to precisely evaluate the contribution of Fc glycan structure to effector functions.

Focusing first on the non-neutralizing antibody CR3022, we employed two complementary assays — ADCC and ADCVI — to characterize Fc-mediated effector functions. Both assays measure antibody-driven activation of effector cells, primarily natural killer (NK) cells, through Fc receptor engagement, leading to the targeted killing of infected cells. However, the two assays measure distinct outcomes. While ADCC assays quantify cytolytic activity against antibody-coated target cells, ADCVI assays assess the ability of antibody-effector cell combinations to suppress viral replication in infected cell populations, a broader functional readout that more closely resembles in vivo antiviral responses. pCR3022 demonstrated significantly greater ADCC activity than mCR3022, as evidenced by a lower EC_₅₀_ value (0.32 µg/mL vs. 27.93 µg/mL, respectively). In the ADCVI assay, pCR3022 effectively reduced infectious viral titers in the presence of PBMCs. This effect was significantly greater than that observed with either mCR3022 or PBMCs alone. These findings demonstrate that Fc-mediated effector functions can endow non-neutralizing antibodies with antiviral activity, and that this activity was strictly Fc-dependent, highlighting the pivotal role of Fc glycoforms in modulating antibody functionality beyond neutralization.

The observed ADCVI activity of pCR3022 against SARS-CoV-2 highlights a potential functional role for non-neutralizing antibodies during infection and raises important implications for vaccine design. In the context of highly mutable viruses like SARS-CoV-2, where escape from neutralization is an ongoing concern, the immune system’s ability to mount Fc-driven responses may represent an underappreciated but critical mechanism of viral control. Unlike neutralization, which requires high-affinity binding to specific viral epitopes, Fc-mediated effector functions rely on the ability of antibodies to recruit and activate immune cells via Fc receptor engagement—a process that is less susceptible to escape mutations [45]. Thus, antibodies that cannot prevent viral entry may still contribute to viral clearance post-infection by facilitating the destruction of infected cells and limiting viral spread. Furthermore, unlike broadly neutralizing antibodies, which are rarely generated during natural infection and are difficult to elicit through vaccination [46,47], non-neutralizing antibodies are commonly induced in nearly all infected individuals, can be readily stimulated by most vaccine platforms, and often persist long-term [48]. Our findings suggest that such antibodies, if equipped with Fc structures that promote effector functions like ADCC, may contribute to early control of SARS-CoV-2 replication—potentially bridging the gap until neutralizing antibody titers are fully developed. This hypothesis is supported by findings in HIV research, where binding antibodies capable of mediating ADCVI appear early in infection and are associated with lower viral loads and improved viral control in both human and animal studies [49,50]. This suggests that similar mechanisms may operate in SARS-CoV-2 infection, where the presence of pre-existing, Fc functionally active, non-neutralizing antibodies—elicited by prior exposure or immunization—could contribute to early control of viral replication, especially during the critical window before neutralizing antibody titers rise to protective levels [51]. Such a mechanism could be especially valuable in countering emerging SARS-CoV-2 variants that escape neutralization by existing antibody therapies. We also propose that future vaccine strategies could benefit from intentionally eliciting ADCVI-inducing antibody responses, providing an additional layer of defense against highly mutable pathogens.

Building on these results, we examined whether the same Fc enhancements could augment the function of a potent neutralizing antibody, CB6. In the ADCVI assay, pCB6 reduced viral titers in a dose-dependent manner in the absence of effector cells, consistent with expected neutralizing activity. However, co-incubation with PBMCs significantly enhanced virus elimination at all concentrations tested, demonstrating a synergistic effect between neutralization and Fc-mediated functions. These findings highlight that Fc effector functions are not limited to compensating for weak or absent neutralization but can also complement and extend the antiviral capabilities of neutralizing antibodies. Thus, incorporating Fc effector functionality into therapeutic antibodies may increase their resilience to viral mutant escape.

Mechanistically, the enhanced ADCC and ADCVI activity observed with GnGn-type glycosylation aligns with prior studies showing that Fc glycan composition critically influences antibody–Fc receptor interactions. Notably, afucosylated IgG antibodies, which those lack core fucose on their Fc N-glycan, exhibit significantly increased affinity for FcγRIIIa, the primary activating Fc receptor on human NK cells [52]. This enhanced binding promotes stronger ADCC responses because FcγRIIIa engagement is essential for initiating cytotoxic effector functions [53]. The GnGn glycoform is inherently afucosylated and contains a uniform, biantennary complex-type structure that is well suited for FcγRIIIa interaction. Our findings are consistent with this model. The GnGn glycovariants of CR3022 exhibited markedly greater ADCC activity compared to its mammalian cell-derived counterpart. These results align with our previous observations that afucosylated glycoforms, particularly the GnGn structure, enhance effector cell activation and further reinforce the functional importance of glycan composition in antiviral immunity [28,54]. Therefore, the improved ADCC and ADCVI activity of our glycoengineered mAbs likely results from more efficient engagement of FcγRIIIa on NK cells and other immune effectors, contributing to their superior antiviral efficacy.

A longstanding concern in antibody-based antiviral therapies, including for SARS-CoV-2, is the potential risk of ADE [55]. To mitigate this, many mAb programs have adopted Fc-engineering strategies—such as LALA mutations—to eliminate Fcγ receptor (FcγR) engagement [56]. While this minimizes ADE risk, it also abolishes beneficial Fc effector functions such as ADCC, as demonstrated in this study. In fact, emerging evidence shows that enhancing Fc functionality does not inherently increase ADE risk. In K18-hACE2 mice, a wild-type anti-RBD mAb provided superior protection compared to its FcγR-null LALA variant, reducing lung viral loads and promoting interferon-γ–mediated antiviral responses [57]. Similar results in hamster and rhesus macaque models confirm that Fc-mediated immunity contributes to in vivo protection without inducing ADE [57]. Our previous work on an anti-dengue mAb showed that the GnGn glycovariant did not promote ADE [58], suggesting the low ADE risk for glycoengineered mAbs such as pCR3022 and pCB6.

In summary, this study highlights the antiviral potential of non-neutralizing antibodies during SARS-CoV-2 infection and the utility of Fc glycoengineering to enhance therapeutic efficacy. Equipping antibodies with uniform GnGn glycans significantly improved ADCC and ADCVI, particularly for the non-neutralizing mAb CR3022. These enhanced Fc functions also synergized with neutralizing activity in CB6, providing a dual mechanism of action. Overall, our findings underscore the importance of Fc effector functions in antibody-mediated protection and the promise of glycoengineering to optimize therapeutic antibodies. Further evaluation of GnGn mAb glycovariants in appropriate animal models will clarify whether the enhanced in vitro activity observed here translates into improved in vivo protection. Moreover, our study suggests that vaccine strategies that elicit both neutralizing and Fc-functional, ADCVI-capable antibodies may offer broader and more durable protection against rapidly evolving viral variants.

## Acknowledgements

We thank Clemens Gruenwald-Gruber and Roman Palt (University of Natural Resources and Life Sciences, Vienna) for performing mass spectrometry analyses. We are also grateful to Dr. Luis Martinez-Sobrido (Texas Biomedical Research Institute) for providing the SARS-CoV-2-delta3a/7b construct. We acknowledge Jennifer Melendez and Daniel Tran for their valuable assistance in maintaining *N. benthamiana* plants.

## Author contributions

Q.C. conceived the study and provided overall supervision. H.Y.S., A.E., H.S., and Q.C. contributed to the experimental design. H.Y.S. and A.E. carried out the experiments and analyzed the data. Q.C. drafted the manuscript with critical revisions from H.Y.S., H.S., and A.E. Q.C. and H.S. also secured funding for the project. All authors have read and approved the final version of the manuscript.

