## Supplemental Figures 1-4, Supplemental Table 1 for "Role of Nonneutralizing Antibodies and Fc Effector Functions in Inhibiting SARS-CoV-2 Infection"

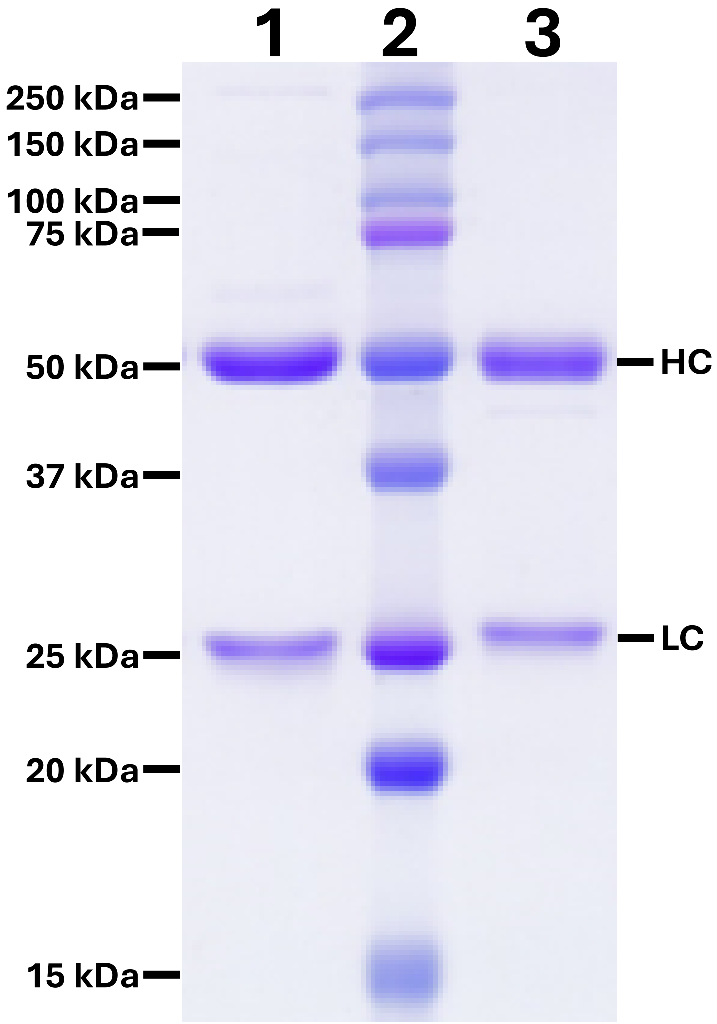


**Figure S1. SDS-PAGE analysis of CR3022 monoclonal antibody expressed in *N. benthamiana* plants.**

Leaves of ΔXFT *N. benthamiana* plants were agroinfiltrated with constructs encoding the heavy and light chains of the CR3022 mAb. Total soluble protein was extracted 7 days post-infiltration, and CR3022 was purified using protein A affinity chromatography. Samples were analyzed by SDS-PAGE under reducing conditions and stained with Coomassie blue. Lane 1: IgG isotype control purified from mammalian cell culture. Lane 2: Molecular weight marker. Lane 3: CR3022 purified from *N. benthamiana*. HC, heavy chain; LC, light chain. Image represents a typical result from multiple independent experiments.

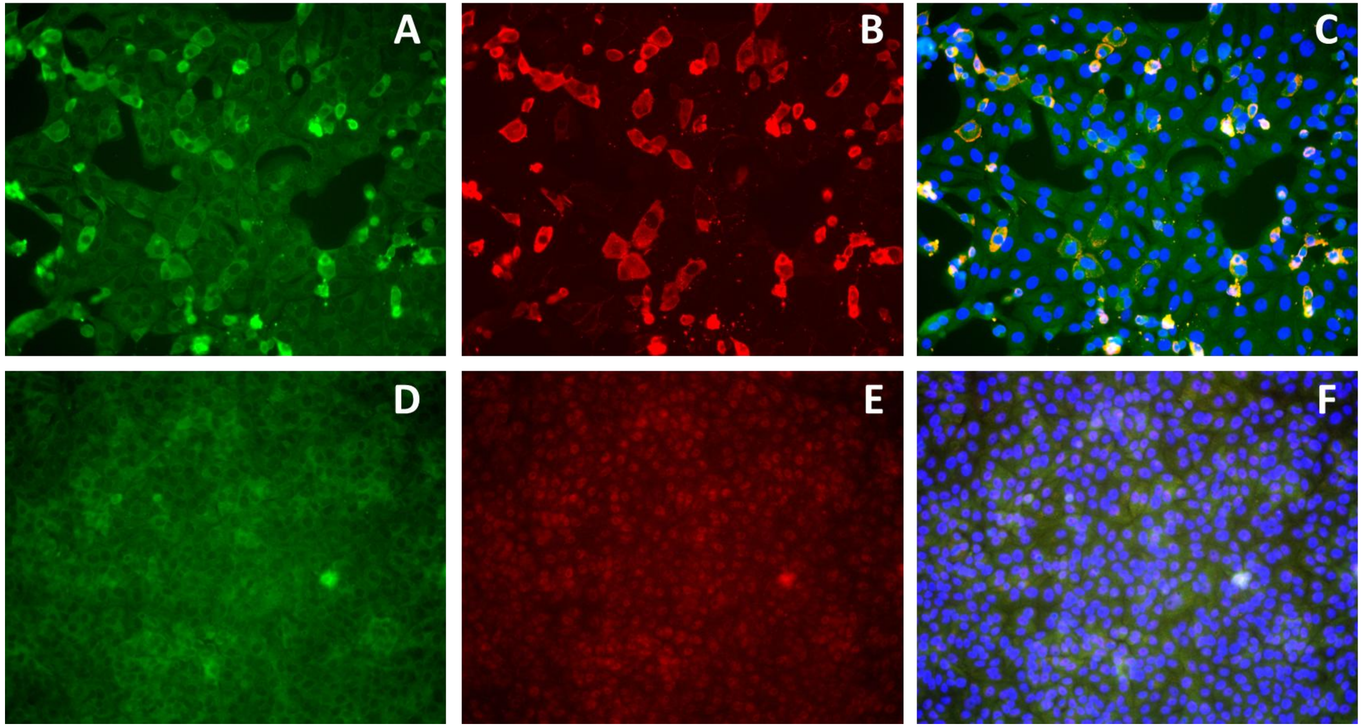


**Figure S2. Recognition of viral antigen in SARS-CoV-2-infected cells by plant-derived CR3022 using immunofluorescence microscopy.**

Vero E6 cells, either infected with SARS-CoV-2 (panels A–C) or uninfected (panels D–F, negative control), were fixed, permeabilized, and incubated with pCR3022 mAb (panels A, D) or an anti-nucleocapsid (N) antibody (panels B, E). Bound antibodies were detected using Alexa Fluor 488–conjugated secondary antibody for pCR3022 (panels A, D) and Alexa Fluor 647–conjugated secondary antibody for the anti-N antibody (panels B, E). Nuclei were counterstained with DAPI (not shown). Merged images of antibody and DAPI staining are presented in panels C (infected) and F (uninfected), corresponding to overlays of A+B and D+E, respectively.

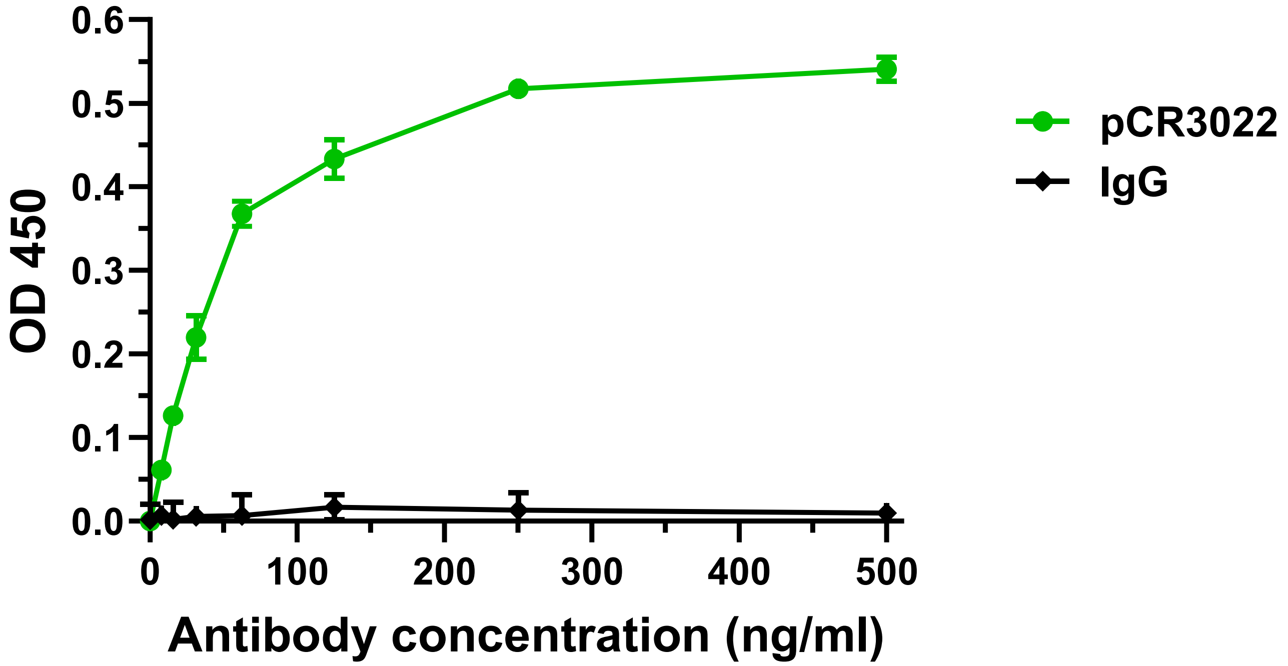


**Figure S3. Specific binding of plant-produced CR3022 to the SARS-CoV-2 receptor-binding domain.**

Serial dilutions of pCR3022 and an isotype IgG control were incubated with immobilized WA1/2020 SARS-CoV-2 RBD on ELISA plates. Binding was detected using an HRP-conjugated secondary antibody, and absorbance was measured at 450 nm. Binding curves and dissociation constants (KD) were generated using GraphPad Prism 10.2.2. Data shown are representative of at least two independent experiments, each performed in technical duplicates. Error bars indicate SD. pCR3022 binding was compared to previously published data for mCR3022 and CB6 [30,33].

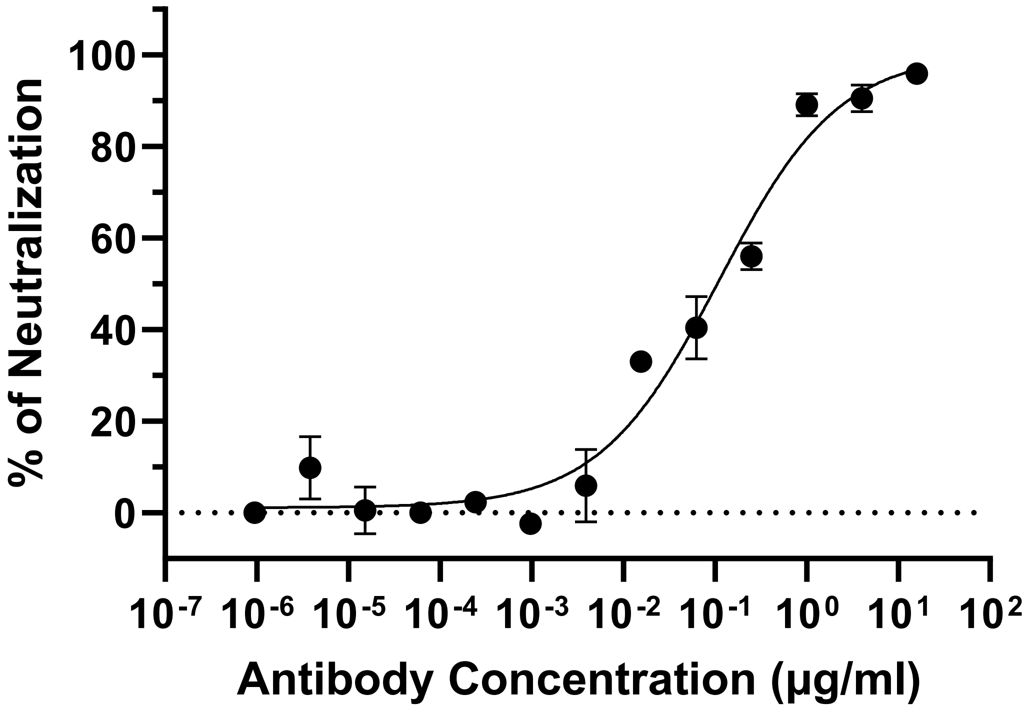


**Figure S4. Neutralization activity of pCB6 against SARS-CoV-2.**

pCB6 was serially diluted and incubated with SARS-CoV-2-delta3a/7b prior to infection of Vero E6 cells. Following infection, cells were fixed, permeabilized, and stained to detect SARS-CoV-2 spike protein. Viral foci were quantified, and percent neutralization was calculated. $EC_{50}$ values were determined using GraphPad Prism (version 10.2.2). Data shown represent at least two independent experiments, each performed in technical triplicates. Error bars represent SD.

| Treatment | Observed Inhibition (%) | HSA Predicted Inhibition (%) | LOEWE Predicted Inhibition (%) | BLISS Predicted Inhibition (%) | ZIP Predicted Inhibition (%) |
|---|---|---|---|---|---|
| **GnGnCB6 ($EC_{75}$) + PBMCs** | **98.51** | **70.62** | **70.33** | **80.99** | **81.41** |

**Table S1. Synergy analysis of pCB6-mediated neutralization combined with effector cell-mediated viral inhibition.**

ADCVI assays were performed using pCB6 alone, PBMCs + pCR3022, or PBMCs + pCB6, with pCB6 and pCR3022 applied at a concentration corresponding to $EC_{75}$ of pCB6. The observed percentage of viral inhibition was recorded for each condition. For the pCB6 + pCR3022 treatment group, predicted inhibition values were calculated based on the PCB6-only and PBMCs + pCR3022 inhibition data using four interaction models available in SynergyFinder.org: Highest Single Agent (HSA), Loewe Additivity (LOEWE), Bliss Independence (BLISS), and Zero Interaction Potency (ZIP), all of which assume no synergistic interaction. Synergy was inferred when the observed viral inhibition from the combination exceeded the model-predicted values, indicating a synergistic effect between antibody neutralization and Fc effector function-mediated viral inhibition.
